# Sensorimotor decisions rely on the entanglement of evidence and motor accumulation processes

**DOI:** 10.1101/2022.05.16.492075

**Authors:** Stijn Verdonck, Tim Loossens, Marios G. Philiastides

**Affiliations:** Faculty of Psychology and Educational Sciences, KU Leuven, Leuven, 3000, Belgium; School of Psychology and Neuroscience, University of Glasgow, Glasgow, G12 8QB, United Kingdom

## Abstract

Most contemporary theories of sensorimotor decision-making formalize the process leading up to a decision as a gradual accumulation of noisy stimulus information over time. The resulting evidence signal is thought to be continuously tested against an internal criterion representing the amount of evidence required to make the decision. In the currently prevailing view, the amount of accumulated evidence required for a decision is independent of the amount of sensory evidence presented by the stimulus, and once that level is reached, a choice is categorically communicated to the motor system to execute an overt response. Recent experimental research casts doubts on both of these assumptions. Using a Leaky Integrating Threshold model, we relax these assumptions specifying both an evidence accumulation and a motor accumulation process. The evidence accumulation signal feeds into a leaky motor accumulator, and it is on the level of the motor accumulation that the final decision criterion is set. This adaptation results in a markedly better description of choice-RT data, especially when it comes to urgency manipulations. We show that this alternative theory, which proposes that sensory evidence is doubly integrated before final evaluation, does not only describe the behavioral data better, but its neural correlates can also be readily derived from EEG signatures involving systems of both evidence and motor accumulation.

## Introduction

The study of sensorimotor decision-making is currently enjoying an ever-increasing amount of attention across different species and levels of neuronal organization (Heekeren et al., 2008; de Lafuente et al., 2015; Hanks et al., 2015; Licata et al., 2017; O’Connell et al., 2018). Central to these endeavors is a computational framework in which noisy sensory information is accumulated over time to an internal decision boundary (Usher & McClelland, 2001; Palmer et al., 2005; Ratcliff & McKoon, 2008). Traditionally, this framework assumed that the decision boundary is independent of the amount of sensory evidence driving the decision and that once the boundary is reached, a choice is categorically communicated to the motor system to execute an overt response.

More recent evidence, however, appears to challenge both of these accounts. For instance, several animal and human electrophysiology studies have reported temporal accumulation profiles terminating at different boundaries, scaling proportionally to the amount of available evidence (i.e., with task difficulty) (Bennur & Gold, 2011; Ding & Gold, 2010; Philiastides et al., 2014; Gherman & Philiastides, 2015; Scott et al., 2017; Herding et al., 2019). Similarly, a wide body of evidence appears to contradict the strict temporal dichotomy between decisional and motor processes and suggests that the (pre)motor system might play a more direct and causal role in decision-making (i.e. embodied cognition) (Filimon et al., 2013; Klein-Flugge & Bestmann, 2012; Thura & Cisek, 2016; Wu et al., 2019; Peixoto et al., 2021; Crapse et al., 2018; Jun et al., 2021).

These recent findings suggest that neural activity related to motor preparation, starts its buildup before the sensory evidence integration completes, effectively lagging the primary process of evidence accumulation (Donner et al., 2009; Servant et al., 2015; Haggard, 2019). Consequently, the amount of sensory evidence could have a direct impact on motor planning, which in turn could influence the eventual choice. This entanglement is not accounted for in current incarnations of decision models nor has it been characterized at the level of neuronal responses (Yau et al., 2021; O’Connell & Kelly, 2021).

Here, we propose an alternative computational framework (Verdonck et al., 2021), which models this entanglement by introducing a secondary, motor-related, leaky integration process that receives the integrated evidence of the primary decision process as a continuous input, and triggers the actual response when it reaches its own threshold. In other words, the primary evidence accumulator relinquishes control of the eventual choice (and hence the strict requirement of an evidence-independent decision boundary) by passing the integrated evidence along to the motor system.

We arbitrate between this alternative theoretical account and conventional single-integration models – including a variant with collapsing decision boundaries (Hawkins et al., 2015; Voskuilen et al., 2016; Palestro et al., 2018; Glickman et al., 2022) – by leveraging computational modeling and time-resolved electroencephalography (EEG) data. We put a special emphasis on a task with a speed/accuracy tradeoff (SAT) manipulation since the SAT offers an intuitive opportunity to exploit the proposed interplay between the primary and motor accumulators.

In doing so, we validate the newly proposed framework and simultaneously demonstrate that the SAT is likely controlled by changes in the proposed leaky motor integration process rather than the long-standing view of boundary adjustments (Ratcliff & McKoon, 2008; Bogacz et al., 2010). Our findings help reconcile the emerging inconsistencies and provide a foundation upon which future studies can continue to interrogate the neural systems underlying rapid sensorimotor decision-making.

## Results

In this work we employed a speeded face-versus-car categorization task (Fig. 1a) (Philiastides et al., 2006, 2011, 2014), while multi-channel EEG data were being collected. The task was designed specifically to induce a time-dependent accumulation of sensory evidence based on concrete perceptual categories through rapidly updating dynamic stimuli (see Materials and Methods for more details). We used two levels of sensory evidence (low and high), by manipulating the phase coherence of the stimuli. We also introduced a SAT manipulation by controlling the amount of time participants had at their disposal to make a response across different blocks of trials (Speed blocks: 1s vs Accuracy blocks: 1.6s). Participants (N=43) indicated their choice via a button press using the index finger of either their right or left hand. The mapping between face/car choices and left/right button presses was counterbalanced across participants. Mean response times were higher in the Accuracy condition (population average of 873 ms) than in the Speed condition (654 ms), while behavioral performance was markedly lower in the Speed condition (population average accuracy of 0.78) compared to the Accuracy condition (0.84) (Fig. 1b; green and black lines, respectively). Finally, for both the Speed and Accuracy conditions, mean response times were higher (by 101 ms, on average) and performance was lower (by 0.15, on average) for the low compared to high evidence trials (Fig. 1b; circles and squares, respectively).

**Figure 1.**
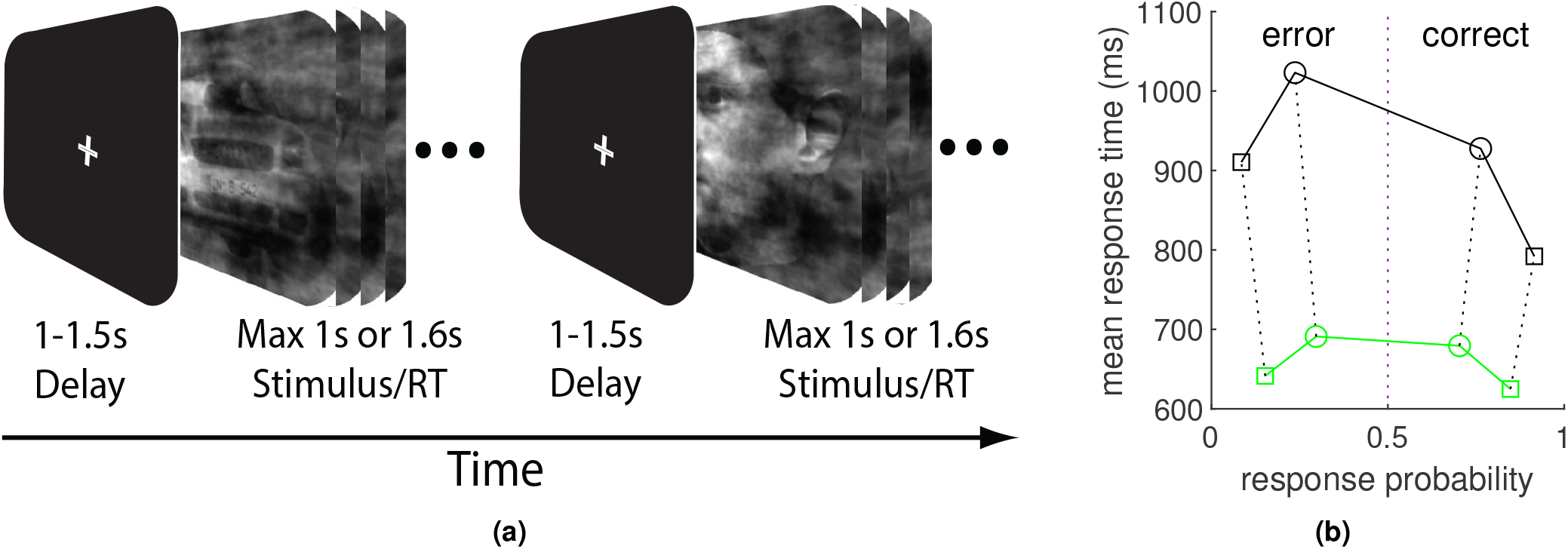
Schematic representation of the experimental task and behavioral performance. (a) Participants had to categorize dynamically updating sequences of noisy images as either a face or a car. In Speed trials participants had up to 1s to decide, while in Accuracy trials they had up to 1.6s. They indicated their choice with either a left or right hand button press, based on a predefined (participant-specific) mapping between the perceptual categories and the two hands. See text for more details. (b) Participant averaged latency-probability plot of face versus car categorization responses, for Accuracy (black line) and Speed trials (green line), respectively. Mean response times are plotted on the left for the error responses (lower probability) and on the right for correct responses (higher probability). Mean response times related to the high evidence condition are denoted by squares and to the low evidence condition by circles.

### The Leaky Integrating Threshold

The current generation of decision making models make a number of fundamental assumptions. Invariably, the presentation of a stimulus is assumed to trigger and drive a process of evidence accumulation in which stimulus evidence is integrated over time. This accumulating evidence is interpreted as a decision variable that typically evolves towards a choice option appropriate for the stimulus that is being presented. A choice is made when the decision variable reaches some criterion or reference state. Finally, this choice is conveyed to the motor system, which in turn delivers the physical response.

In Verdonck et al. (2021), we break with this tradition and propose to broadcast the accumulated evidence into a secondary, leaky motor accumulator and assume that the stopping criterion applies only for this secondary stage of the decision making process. In other words, there is no longer a binary choice stemming from the initial evidence accumulation which then triggers a motor response but rather from a secondary motor accumulation process that simultaneously accumulates to its own threshold by taking the evidence emerging from the first process as input. This ‘Leaky Integrating Threshold’ (LIT) effectively entangles evidence and motor accumulation, resulting in a host of specific predictions that can be falsified using both behavioral and neural data. The LIT can be applied to any model of decision making, but in this paper we use it to extend the common constant drift diffusion model (DDM).

Mathematically, the dynamics of the deterministic LIT are defined as

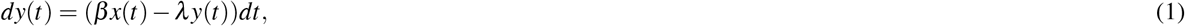

with *x*(*t*) the evidence accumulated over time and *y*(*t*) the resulting motor accumulation. For the analyses in this paper we will use a simple constant drift diffusion process for the dynamics of *x*(*t*) with stimulus dependent drift speeds *v_i_* (*i* denoting the stimulus condition). The boundaries are not imposed on the accumulating evidence *x*(*t*) directly, but on the emergent motor accumulation *y*(*t*): we denote the characterizing boundary separation with *a*, with an upper boundary at 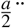 and a lower boundary at 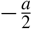. We assume the starting position of the evidence accumulation *x*(0) = *x*_0_. The leak parameter *λ* determines the lag between the evidence and motor accumulation and *β* is a scaling parameter. When the motor accumulation in *y*(*t*) hits a threshold, the physical response ensues and a choice is made. In terms of choice RT data alone, the LIT has one redundant parameter: to make the model estimable we choose *β* = *λ*. Solving for *y*(*t*) in Equation 1 results in a time smoothed version of the original accumulating evidence:

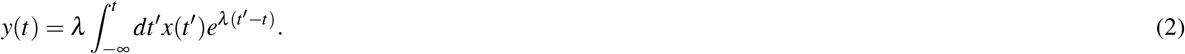

The time smoothing afforded by the motor accumulation effectively increases the signal-to-noise ratio of the primary evidence accumulation signal and reduces accidental noise-driven threshold crossings. Unlike traditional instances of the DDM in which the momentary state of the accumulated evidence itself is being evaluated, here decisions are based on a time-decaying weighted memory of all the evidence accumulator’s past values (hence the term Leaky Integrating Threshold). This introduces a lag between the primary evidence accumulation and motor accumulation processes. The leak parameter *λ* determines the extent of the time smoothing and the size of the corresponding lag. When *λ* approaches infinity, the LIT has no memory or lag and behaves as a normal threshold on the primary accumulated evidence. Assuming pre-stimulus evidence accumulation is *x*(*t* < 0) = *x*_0_ and motor accumulation *y*(*t*) has had enough time to adapt, the motor accumulation starting position becomes *y*(0) = *x*_0_.

### Leak vs boundary adjustments for controlling choice urgency

To manipulate the urgency to make a response, participants completed four blocks of the face-vs-car categorization task either under a Speed or an Accuracy instruction (Fig. 1a). To fit the behavioral data (Fig. 1b) we proceeded with the estimation of the LIT as follows: First, for the evaluation of its choice-RT probability density function one needs to resort to time consuming numerical integration (e.g., simulations). We deal with this computational bottleneck by using the prepaid method proposed by Mestdagh et al. (2019). Second, in order not to impose any constraints on the shape of the non-decision time distribution, we opt for a D*M fit criterion (Verdonck & Tuerlinckx, 2016). For this analysis, Speed and Accuracy data are fitted completely independently, resulting in different parameter values for each instruction. In line with Verdonck et al. (2021), we found that inverse leak *λ*^−1^ and boundary separation *a* were significantly smaller in the Speed than in the Accuracy condition looking at their intra-individual differences (Δ*λ*^−1^ = −0.19 [−0.22, −0.15], Δ*a* = −0.089 [−0.12, −0.061]). In addition, the relative starting position *zr* and drift speeds *v*_*i*=1…4_ were indistinguishable (Δ*zr* = 0.016 [-0.024,0.059], Δ*v*_1_ = −0.044 [−0.24,0.14], Δ*v*_2_ = −0.27 [−0.79,0.033], Δ*v*_3_ = −0.085 [-0.29,0.11], Δ*v*_4_ =0.19 [-0.15,0.6]). The bracketed numbers are the 99% bootstrap confidence intervals.

While the prepaid method is useful for the estimation of parameters of models without a readily accessible likelihood function, it does not allow a direct comparison between models. For model comparison, we use an approximation of the LIT, as has been introduced in Verdonck et al. (2021) and further developed in Materials and Methods below. This approximation is a reparameterized version of a standard diffusion model, in which the original LIT leak parameter translates to a change in the boundary separation, starting position and non-decision time (Equations 11, 4 and 3). This allows us to construct an overarching model in which multiple mechanisms of controlling for choice urgency can be compared (Equation 12). Specifically, we compare the LIT based mechanism (balancing the signal-to-noise ratio of a secondary motor accumulation to bound by controlling its leak parameter) with the two most common alternatives: controlling a single stage evidence accumulator’s threshold, either by changing its overall value (Ratcliff & McKoon, 2008; Bogacz et al., 2010), or allowing different rates of boundary collapse (Hawkins et al., 2015).

Because the dynamics of this model set are described by a simple one-dimensional diffusion process, we can use a grid method as in Voss & Voss (2008) to calculate the associated likelihood function. Again using D*M as a fit criterion, we estimate three implementations, each of them relying on a different parameter to explain the differences between urgency conditions in the data: inverse leak (*λ*^−1^), boundary separation (*a*), or collapse rate (*c*). As all of the resulting models have the same number of free parameters, we can use their fit to the data as a way to arbitrate between the mechanisms (Fig. 2). For the great majority of participants, the LIT based inverse leak parameter (*λ*^−1^) is more successful at explaining differences between urgency conditions than both simple boundary separation *a* (72%[51%, 88%]) and collapse rate *c* (81%[63%, 93%]). The bracketed numbers are the 99% bootstrap confidence intervals.

**Figure 2.**
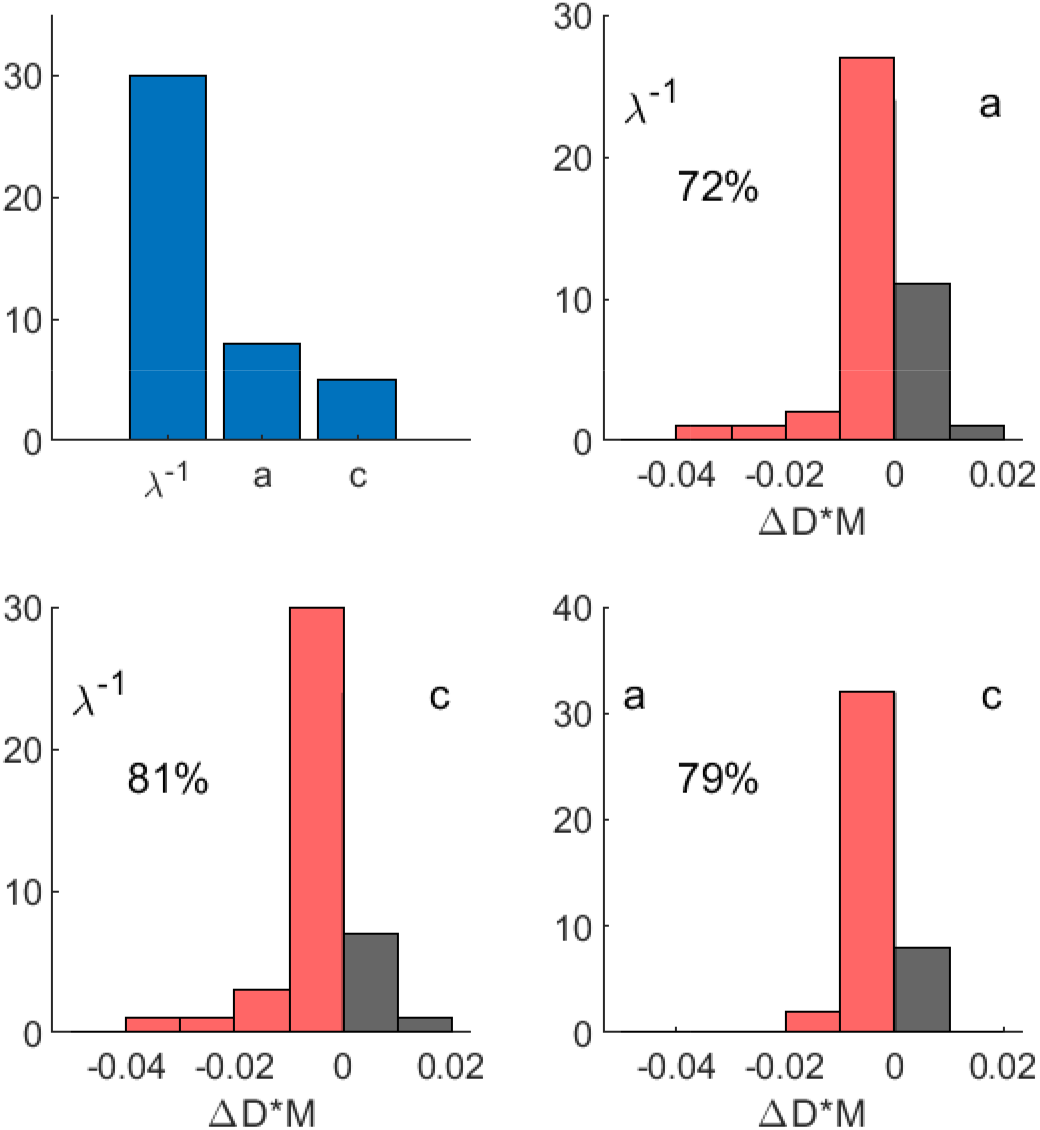
Comparison of 3 competing mechanisms for controlling choice urgency. In the top panel, the amount of participants favoring each of the 3 mechanisms. Each of the 3 plots below show the distribution of differences in fit Δ*D* * *M* between every combination. Additionally, the percentage of Δ*D* * *M* below zero is displayed, indicating how frequently the preferred mechanism was chosen in favor of the respective alternative.

### Electrophysiological predictions of LIT versus standard DDM

To connect any high level model of evidence accumulation to EEG data, assumptions have to be made about how exactly the theoretical signal can be expected to manifest at the level of macroscale brain dynamics. In accordance with previous work (Kelly & O’Connell, 2013a; Philiastides et al., 2014), we assume that accumulation activity on the scalp (e.g., gradual build-up of the EEG signal) will be observed as having the same sign for both choice alternatives. To accommodate this in a framework of one dimensional evidence accumulation, the theoretical evidence signal can be recast as an absolute deviation of a pre-stimulus baseline.

Moreover, as EEG data could reflect a superposition of different processes, we cannot assume that a clean representation of the evidence accumulation signal can be found at any particular location or for any combination of sensors. Regardless of the unmixing technique used (Parra et al., 2005), the evidence accumulation signal extracted from the EEG data might contain activity from other processes. If we start from the premise that a meaningful model of decision making will at least include processes that explain some of the differences in choice RT between trials, a valid comparison between model-derived and actual EEG accumulation signals could consist of looking at inter-trial correlations of momentary signal values with their respective final RTs, instead of comparing the raw accumulation signal profiles themselves.

This approach has the important advantage that EEG activity that is shared across trials but does not influence final choice RT outcomes (and hence may be unrelated to the process of interest) is filtered out. More specifically, correlations of momentary signal values with their corresponding final RTs are not affected by an ‘unspecific’ shift in the signal values, as long as it is present in all trials. In other words, adding a constant signal to all trials - although very much altering the resulting mean accumulation signal - would not impact the correlation of momentary accumulation values and RTs. These “accumulation-RT” correlations can be calculated in a response-locked fashion, for different points in time leading up to the eventual response.

The ensuing accumulation-RT correlation curves have an amplitude that can be interpreted as the extent to which eventual trial RTs can be predicted by their preceding accumulation signal at a given point in time. The sign of the accumulation-RT correlation indicates whether higher accumulation signals lead to higher RTs or the other way around. In trials where the accumulation signals of slow and fast trials are comparable in value, the accumulation-RT correlation is zero. On the whole, accumulation-RT correlation curves are driven by how different trials (from slow to fast) evolve compared to each other, and not how their mean value evolves compared to the rest of the brain activity. Accumulation-RT correlation curves can therefore be seen as a robust metric to compare the internal dynamics of theoretical evidence and motor accumulation signals with what is observed in the EEG signals.

To assess these properties in both the standard DDM and its LIT extension, we simulated all accumulation signals and their corresponding accumulation-RT correlations. In Figure 3a, participant averaged DDM evidence accumulation signals for two levels of sensory evidence are shown locked to the motor response. At all times before the response, the low sensory evidence signal is higher than the high sensory evidence signal (i.e., the accumulation signals never cross before the final response). The difference between the two signals reduces to zero only at *t* = −∞ and at the response (*t* = 0). The latter is consistent with the idea of a stimulus invariant information threshold, whereby all evidence accumulation signals at *t* = 0 have the same value. Correspondingly, the evidence accumulation-RT correlation curve is always positive (higher values lead to higher RTs) but reduces to zero at *t* = −∞ and *t* = 0.

**Figure 3.**
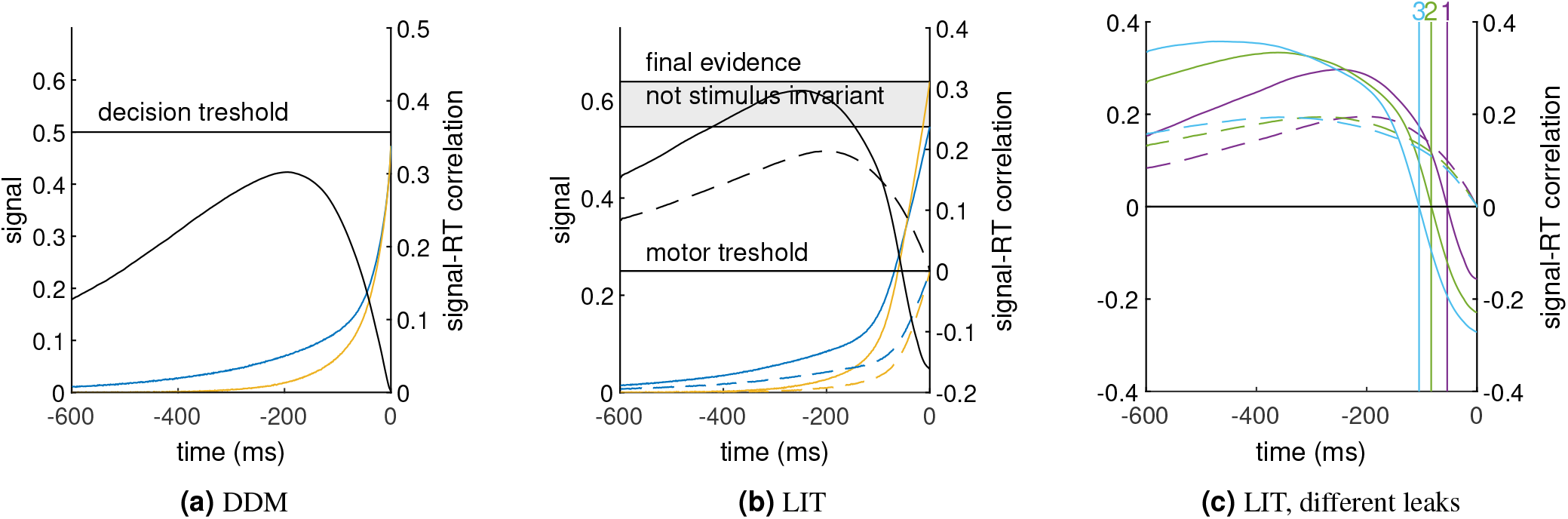
Model predictions. (a) Mean evidence accumulation signals of a DDM, response locked, for two different levels of sensory evidence (low in blue, high in yellow): their values correspond to the left vertical axis. In grey and corresponding to the right axis, is the momentary correlation of all individual trials with their final response times, calculated for every response locked time before the actual response. (b) Mean evidence and motor accumulation signal of a LIT for two levels of sensory evidence (low in blue, high in yellow; solid lines represent evidence accumulation, dashed lines motor accumulation) and the corresponding momentary correlations with response times in grey. (c) Momentary correlations with response times of decision (solid) and corresponding motor accumulation signals (dashed) for thee different values of the leak parameter. The zero correlation crossings of the respective decision signals are indicated with vertical lines, annotated with the index of the leak parameter, as can be found in Table 2 in Materials and Methods.

This observation stands in stark contrast with the behavior of the LIT (Fig. 3b) and recent EEG work demonstrating a stimulus-depended signal modulation at the time of response (Philiastides et al., 2014; Herding et al., 2019; von Lautz et al., 2019). The mean LIT evidence accumulation signal for low sensory evidence dominates the signal for high sensory evidence at the early stages, but later the two signals cross over as the signal for high sensory evidence begins to dominate. At the response (*t* = 0), the signals for the two levels of sensory evidence do not coincide, demonstrating a clear deviation from the idea of a stimulus invariant information threshold on the level of the evidence accumulation. This pattern also manifests in the accumulation-RT correlation curve via a sign switch at the cross-over point between the evidence accumulation signals, ending up negative at the time of response.

For the LIT the final decision criterion is not acting on the evidence accumulation itself, but rather on a secondary motor accumulation (Fig. 3b; dashed lines) that is driven by the primary evidence accumulation. The corresponding motor accumulation-RT correlation curve equals zero at the response, consistent with the fixed criterion imposed on this accumulator. Finally, we demonstrate how the LIT leak parameter impacts the profile of the accumulation-RT correlation curves (Fig. 3c). Specifically, higher values of inverse leak *λ*^−1^ (i.e., more time smoothing applied going from evidence accumulation to motor accumulation) result in an increased lag between motor and evidence accumulation signals. This lag can be quantified by the response-locked time at which the accumulation-RT correlation curve crosses zero (Fig. 3c; vertical lines).

In the next section, we compare these theoretical accumulation-RT curves with the corresponding curves derived from EEG accumulation signals to offer neurobiological validation for the proposed LIT framework.

### EEG signatures of LIT-like evidence accumulation

To offer neurobiological validation for the LIT we first aimed to identify candidate accumulator signals from which to derive the required accumulation-RT curves. Specifically, we used a single-trial multivariate classifier (Diaz et al., 2017; Franzen et al., 2020; Philiastides & Sajda, 2006) designed to estimate linear spatial weightings of the EEG sensors discriminating between low and high levels of sensory evidence (collapsing across Speed and Accuracy trials) as was done in Philiastides et al. (2014) (see Materials and Methods). Applying the estimated electrode weights to single-trial data produced a measurement of the discriminating component amplitudes, which we treat as a neural surrogate of the relevant decision variable being integrated. We performed this analysis on response-locked data and separately for each participant.

Importantly, this approach offered an initial opportunity to arbitrate between the predictions of the traditional DDM and the LIT. The traditional DDM assumes a common threshold for the primary evidence accumulator for both low and high evidence trials. This in turn should lead to low discrimination performance near the response with systematic improvements in classification accuracy further back in time as the traces associated with the different levels of sensory evidence begin to deviate (Fig. 3a). In contrast, the LIT predicts that classification accuracy, in so far as it is driven by the evidence accumulation signal, should be highest near the response, drop gradually moving back in time as the traces of the different levels of sensory evidence cross and pick up again as they begin to deviate in the opposite direction (Fig. 3b). In other words, we should observe two separate discrimination peaks in the period leading up to the response.

We note that the LIT motor accumulation signal (i.e., secondary accumulation) may also contribute to the classification accuracy further away from the response, but, because it converges to a common threshold, cannot explain any potential above chance classification near the response itself. We observed two local discrimination maxima, one near the response itself and one approximately 400 ms prior the response, on average (Fig. 4a). This classification performance profile fits squarely within the LIT predictions, offering initial support for this framework. The spatial topography (forward model; see Materials and Methods) of the discriminating activity at the time of response (Fig. 4b) is consistent with the spatial distribution of centroparietal EEG signals implicated previously in the process of evidence accumulation (O’Connell et al., 2012; Kelly & O’Connell, 2013a; Philiastides et al., 2014).

**Figure 4.**
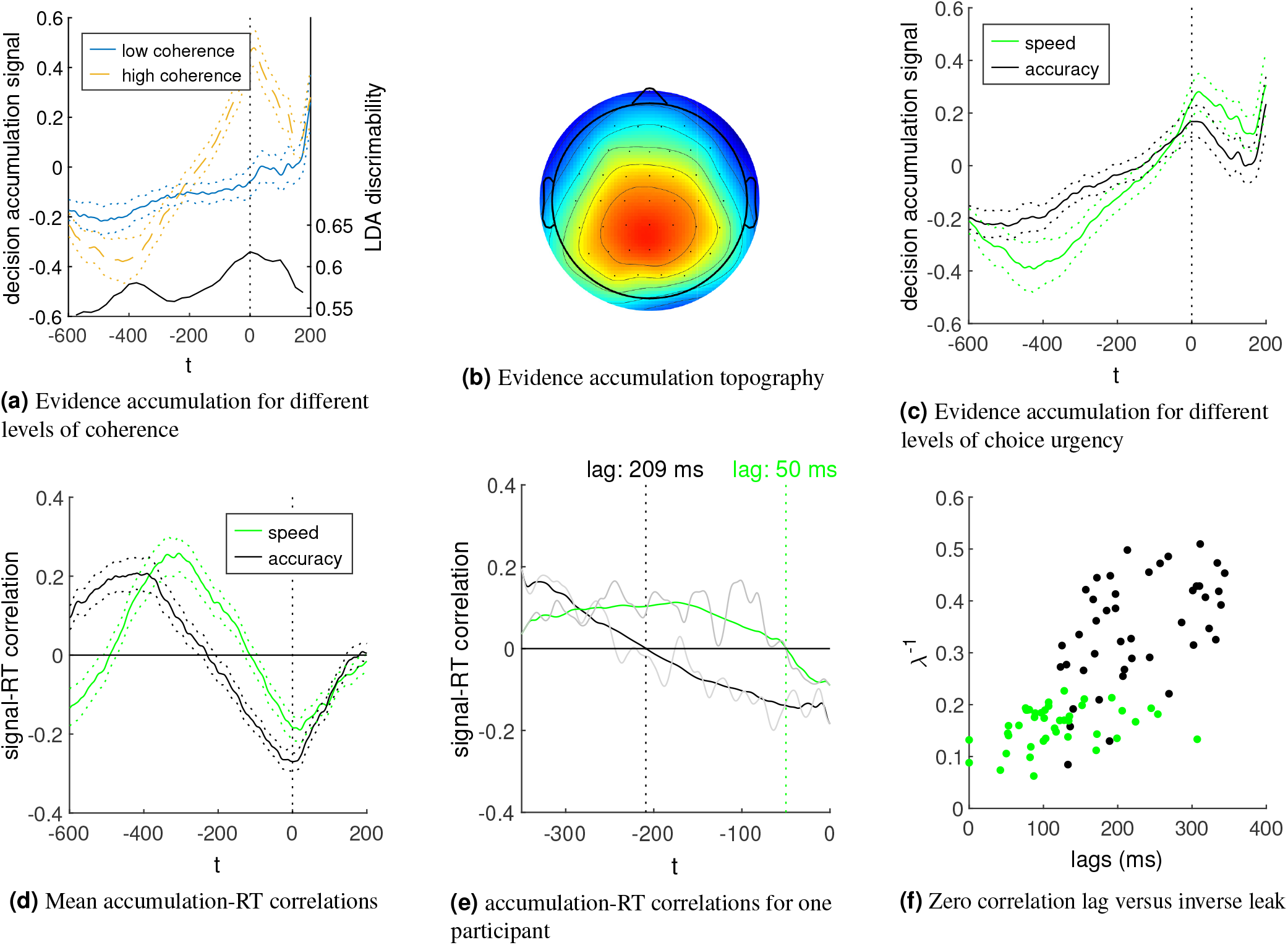
Evidence accumulation signals in the EEG. (a) Participant average evidence accumulation signals for low and high sensory evidence. (b) Participant average scalp topography (forward model) associated with the traces in (a). (c) Participant average evidence accumulation signals for speed and accuracy conditions. (d) Participant average accumulation-RT correlation curves derived from the momentary inter-trial signal correlations with final RTs for speed and accuracy conditions. (e) Accumulation-RT correlation curves of a single participant in the speed and the accuracy condition, with corresponding motor-lags indicated by the dashed vertical lines. (f) Scatter plot of each participant’s motor-lag (EEG-derived) with their inverse leak (LIT model-derived). All error bars represent 95% confidence bounds on the corresponding signals.

We subsequently applied each participant’s spatial weights obtained in the 50ms time-window centered around the response (i.e., window of best overall discrimination performance) on the entire response-locked time window to obtain individual trial-by-trial evidence accumulation signals. This approach can be thought of as projecting the data through the same neural generators responsible for the main discriminating activity and was designed to obtain robust estimates of the full temporal profile of this activity beyond the point of maximum discrimination (Philiastides et al., 2014; Ratcliff et al., 2009).

An initial qualitative assessment of the resulting temporal profiles is in line with what is predicted by the LIT (Fig. 3b), whereby there is no common decision boundary for low and high sensory evidence trials close to the response. On the contrary, it is the moment where the difference between the two signals is most pronounced (Fig. 4b). Additionally, the signals appear to cross over some time before the response (circa 200 ms prior to the response), creating two regions where they are separable: an early one where the high sensory evidence signal is lower than the low sensory evidence signal, and a later one near the response itself where the effect is reversed. Similarly, stratifying trials according to the SAT manipulation suggests there are no boundary differences across the speed and accuracy evidence accumulation signals near the response, on average 4c).

Importantly, these temporally-resolved evidence accumulation signals could be examined formally in the framework of accumulation-RT correlation curves and be directly compared to the theoretical predictions of the LIT. This approach is generally more robust against potentially interfering trends from unspecific neural processes that mix into the EEG signal and are not coupled to RT. In line with the LIT model predictions, both the Speed and Accuracy evidence accumulation signals show trial-by-trial correlations with RT that are negative on response and become increasingly less negative going back in time until they cross the zero correlation line while they continue to become increasingly positive (i.e., compare Fig. 3c and 4d).

Next, we define motor-lag as the time between the zero-crossing of the evidence accumulation-RT correlation curve and the actual response, since we expect the motor accumulation signal to reach zero correlation with RT very close to the response itself (this is implied by a fixed threshold, where the signal values of all trials are equal). If we assume the inverse leak parameter *λ*^−1^ in the LIT is largely responsible for SAT control (as shown in Verdonck et al. (2021) and replicated here – the inverse leak is larger for accuracy than speed in 42 out of 43 participants), then the LIT makes a very specific prediction: the motor-lag for Speed should be markedly smaller than the one for Accuracy (Figure 3c). We indeed see this pattern emerge in our population EEG data (Fig. 4d) as well as in individual participants (Fig. 4e).

Finally, we also find a clear correlation between *λ*^−1^ (obtained solely from behavioral choice-RT data) and the individual motor-lags (obtained purely from the EEG data), such that participants with higher inverse leak *λ*^−1^ also exhibit a higher motor-lag (a percentage bend correlation of 0.68 with a 95% CI of [0.55,0.8]; Fig. 4f). This is consistent with the notion that higher values of *λ*^−1^ lead to more time smoothing for the motor accumulation, thus increasing its signal-to-noise ratio and decreasing the chance of responding incorrectly, while at the same time leading to an increase in motor-lag, resulting in slower responses.

### EEG signatures of LIT-like motor accumulation

Having established the presence of evidence accumulation signals in the EEG consistent with the LIT, we then tested the extent to which the relevant motor accumulation signals could also be captured at the macroscopic level of scalp potentials. Theoretically, the motor accumulation signal can be derived from an already established evidence accumulation signal. Specifically, Equation 1 suggests there should be a linear relation between the change of the motor accumulation signal (i.e., slope) and the value of the evidence accumulation signal, at every time *t* the LIT is operational. We applied this theoretical framework to derive motor accumulation signals directly from the EEG signals, which we then used as a benchmark for further validating the LIT framework.

More concretely we first calculated, for each trial separately, the local raw signal slopes for all EEG channels (in windows of 50 ms in duration) at different times *t*. As detailed in Materials and Methods, we calculated these slope snapshots (each consisting of 64 slope values - one for each channel) every 25 ms, with window centres ranging from 150 ms to 50 ms before the response. We expect the build-up of the proposed motor accumulation signals to be most pronounced in this interval due to increases in corticospinal excitability in anticipation of the eventual motor response (Chen et al., 1998). Similarly, this is the interval over which evidence accumulation signals peaked (see Fig. 4a) and hence the leaky integrating response of the motor accumulator should be most pronounced during this period.

From the slope snapshots (# trials x # time windows), we then estimate, for every participant separately, the optimal sensor weights for maximising the correlation between the momentary slopes in the raw sensor signals and the concurrent values of the evidence accumulation signals we derived in the previous section (see Materials and Methods). According to the LIT, the momentary slope of a motor accumulation signal should indeed be proportionate to the corresponding momentary value of evidence accumulation (Equation 1). By finding the weights that make this feature most pronounced, we construct a motor accumulation signal that is maximally compatible with one of the fundamental assumptions of LIT dynamics. We calculated these weights using only the trials of the high sensory evidence condition because these trials have the most pronounced evidence accumulation signals and will be better at teasing out the corresponding motor accumulation slopes.

Next, we applied each participant’s resulting sensor weights on the entire response-locked time window to obtain robust estimates of the full temporal profile of individual trial-by-trial motor accumulation signals (Fig. 5a). Furthermore, we estimated the spatial topography of the proposed motor accumulating signal by calculating its correlation with each individual sensor signal, using data from all trials (Fig. 5b) as well as separately for left and right choice trials (Fig. 5e, f) in order to harvest the well-known contralateral motor bias (Donner et al., 2007, 2009). Importantly, these topographies are calculated purely based on LIT model assumptions and EEG-derived evidence accumulation signals. As such they are not informed about the laterality of the motor response, which we will use as further validation for the presence of motor accumulation signals.

**Figure 5.**
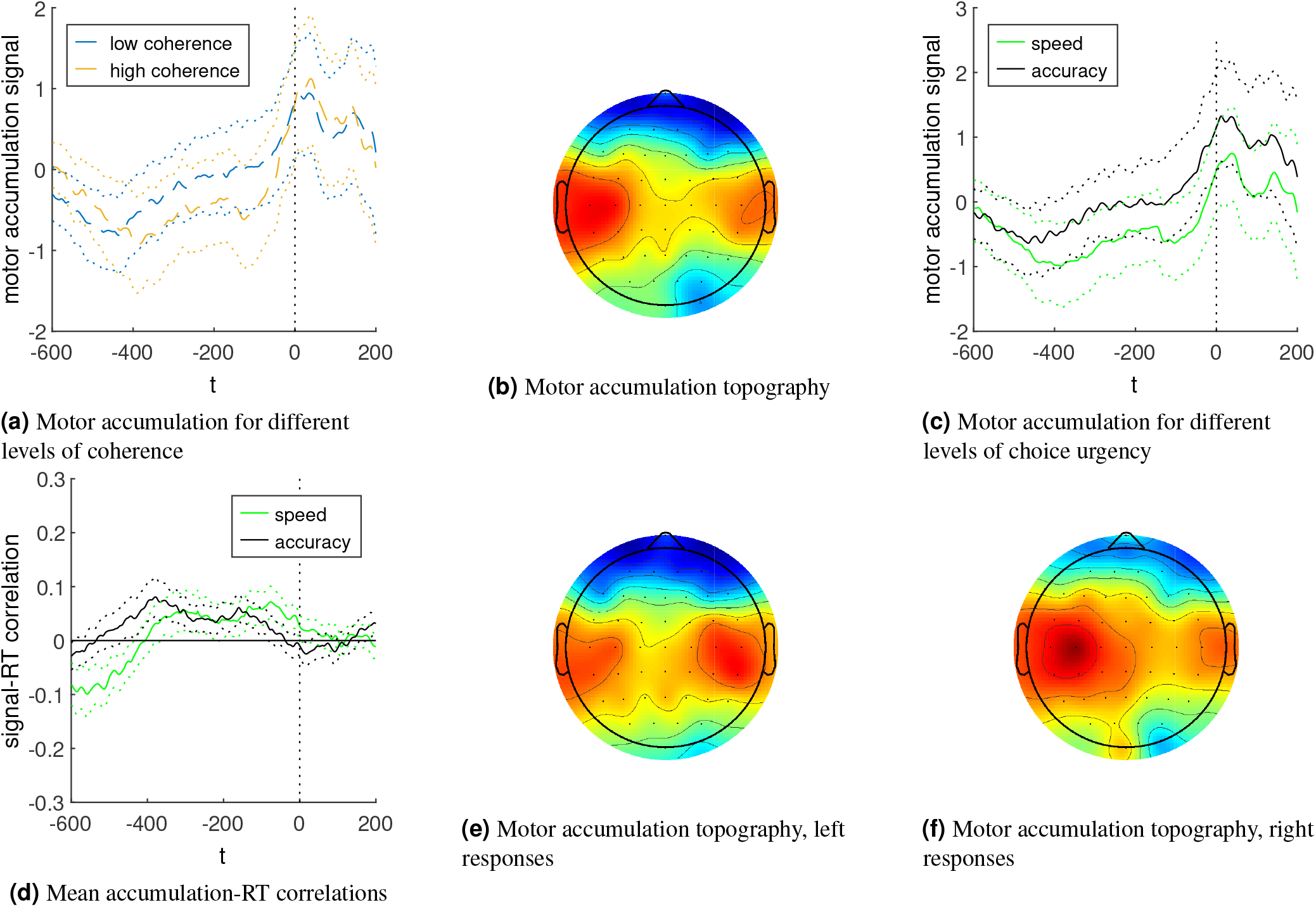
Motor accumulation signals in the EEG. (a) LIT-informed motor accumulation signals for low and high sensory evidence trials, averaged across participants. (b) Participant average scalp topography associated with the traces in (a) based on their across trial correlations with the observed EEG signals at each sensor at response time. (c) Participant average evidence accumulation signals for speed and accuracy conditions. (d) Participant average accumulation-RT correlation curves derived from the momentary inter-trial signal correlations with final RTs for speed and accuracy conditions. (e, f) Participant average scalp topographies for left and right hand responses, respectively. All error bars represent 95% confidence bounds on the corresponding signals.

The temporal profile of the proposed motor accumulation signal exhibits a gradual build-up of activity, which peaks at the time of response (Fig. 5a,c). Given the high uncertainties (dotted curves) of the raw signal estimates, it is difficult to judge differences between low and high sensory evidence trials (Fig. 5a), or Speed and Accuracy trials (Fig. 5c). Additionally, these estimates could be contaminated by signals that show differences between conditions but have no relation to the actual motor accumulation process. Once again, we exploit the inter-trial accumulation-RT correlation curves to circumvent some of these potential contaminants to validate the specific temporal features of the LIT motor accumulation signal.

Comparing the trial-by-trial accumulation-RT correlation curves between Speed and Accuracy conditions (Fig. 5d), we observed strong agreement with the LIT predictions (Fig. 3c). Specifically, the Speed trials (high integration leak) have their maximum accumulation-RT correlation closer to the response compared to Accuracy trials, and both conditions end up close to a zero accumulation-RT correlation around the response. These observations further reinforce the notion of a common motor accumulation boundary and a SAT implementation via leak adjustments of the secondary, motor accumulator.

Finally, we used the spatial topographies of the process of motor accumulation to offer further validation that these signals are indeed emerging from (pre)motor brain structures over the motor strip. Motor preparatory activity has a prototypical scalp distribution with activation clusters over centrolateral sensors and a pronounced contralateral response bias. Here we mapped the two choices on separate hands and we would therefore expect scalp topographies associated with motor accumulation signals to peak contralaterally to the motor effector used to indicate each choice (i.e., over left [right] centrolateral sensors for right [left] button presses). The topographies we derived for our motor accumulation signals are perfectly consistent with the aforementioned scalp profiles (Fig. 5b, c), offering further validation of the LIT.

## Discussion

In this work we propose a new computational framework for operationalizing sensorimotor decisions (Verdonck et al., 2021) and offer neurobiological validation of the proposed mechanism using human electroencephalography data. Specifically, in the context of traditional evidence accumulation models of choice-RT we introduce a secondary motor-related leaky integration process (LIT), which receives already stochastically accumulated evidence from a primary process of evidence integration. This doubly integrated signal manifests as a lagged and smoothed version of the original evidence accumulation signal and controls the eventual choice as it crosses its own threshold, while allowing the primary accumulation process to remain unbounded.

This framework was borne out of necessity to reconcile discrepancies between traditional diffusion models and recent experimental observations in the literature. Firstly, there has been evidence pointing to stimulus-depended values of accumulated evidence at the time of choice (contrary to the traditional view of a common decision boundary) (Bennur & Gold, 2011; Ding & Gold, 2010; Philiastides et al., 2014; Gherman & Philiastides, 2015; Scott et al., 2017; Herding et al., 2019). Secondly, there has been support for a more active role of (pre)motor areas during sensorimotor decisions (Filimon et al., 2013; Klein-Flugge & Bestmann, 2012; Donner et al., 2009; Thura & Cisek, 2016), with inactivation of such regions leading to gross behavioral impairments (Wu et al., 2019; Peixoto et al., 2021; Jun et al., 2021; Erlich et al., 2015; Crapse et al., 2018).

An emerging trend in the literature focuses on the potential utility of single-accumulator models with collapsing boundaries (Hawkins et al., 2015; Voskuilen et al., 2016; Palestro et al., 2018; Glickman et al., 2022). This class of models, incorporates a continuously encroaching deadline, which dynamically changes the evidence criterion to account for the urgency to make a response. Here, we offer an alternative formulation that additionally considers the likely role of the motor system, while at the same time offering a more flexible account of decision making with potentially wider implications (see below).

In the case of collapsing boundaries, the model adjusts the end criterion by monitoring the noisy accumulated evidence at discrete points in time without “remembering” past observations (i.e., boundaries are “memoryless”). In the LIT, instead of actively changing the end criterion of the primary accumulator, a second (motor) integration is used to make the signal less noisy to avoid unwanted accidental crossings. In this framework, the motor boundary has a leaky memory, which enables it to consider past evidence, rather than just monitor the integrated evidence for an immediate crossing.

Here, we first show computationally that the LIT not only offers a better fit to behavior compared to traditional single-accumulator diffusion models, but it also provides a vitally different perspective on how choice urgency alters decision dynamics. Contrary to the contemporary view that boundary adjustments (dynamic or otherwise) control the speed versus the accuracy of a decision (Ratcliff & McKoon, 2008; Bogacz et al., 2010), here we demonstrate that it is a change in the leak of the motor accumulator that instantiates this tradeoff. These findings also help reconcile recent neural and computational reports showing little evidence of collapsing decision boundaries during SAT manipulations (Heitz & Schall, 2012; Hanks et al., 2014; Voss et al., 2019; Rafiei & Rahnev, 2021).

Next, we offer neurobiological validation of the proposed LIT dynamics by leveraging high temporal precision human electrophysiological signals against which to compare the relevant modeling predictions. We capitalize on the fundamentally different accumulation signal predictions of the LIT compared to traditional diffusion models to arbitrate between the competing theoretical constructs. Specifically, we focus on how EEG accumulation signals manifest spatially on the scalp, how they behave as the decision unfolds as a function of stimulus-evidence as well as how they dynamically relate to the eventual choice RTs.

In line with the LIT predictions, we identify separate signatures of decision-related accumulation with two distinct spatiotemporal profiles. Specifically, an initial evidence accumulation signal with a centroparietal activation profile (Kelly & O’Connell, 2013b; Philiastides et al., 2014; van Vugt et al., 2019; Luyckx et al., 2020) is trailed by a secondary accumulation signal emerging over the motor strip, with higher activity contralateral to the motor effector used to indicate the decision, consistent with the well-known contralateral motor bias (Donner et al., 2007, 2009). Importantly, we also show that the temporal lag between the two EEG accumulation signals across participants scales systematically with the model-derived rate of information leakage in the motor accumulator. To further alleviate potential confounds related to the quality of the unmixing of the two EEG accumulation signals we also look at the inter-trial (i.e., point-wise) correlations of these signals with their final RT outcomes and are able to reliably reproduce their theoretically-derived counterparts.

To our knowledge, this is the first instance of a model in which a secondary motor accumulation process becomes part of the causal chain of events and drives the decision after receiving stochastically accumulated evidence from a primary source of evidence integration. We believe this mechanistic proposition is the main novelty of this work, which could have wider implications in how sensorimotor decisions are being formed. For example, the LIT could be relevant in scenarios involving integration of non-stationary evidence with varying degrees of temporal uncertainty (Ossmy et al., 2013) by controlling the integration time constant and adjusting the leakage of information being used by the motor accumulator.

An additional advantage of the LIT over previous work is that choice is determined by the motor accumulator, which allows the primary accumulator to remain unbounded and continue to accumulate past the decision. This in turn could potentially inform secondary decisions involving additional post-decisional deliberation (e.g. “change-of-mind” decisions, post-decision metacognitive appraisal, etc) (van den Berg et al., 2016; Resulaj et al., 2009; Pleskac & Busemeyer, 2010) or even influence information processing on subsequent choices (i.e., introduce serial dependencies across trials) (Urai et al., 2019; Desender et al., 2019).

For example, consider the case of change-of-mind decisions or double responses (Resulaj et al., 2009; Evans et al., 2020). Because the LIT is a separate motor accumulator, resetting it after a motor action does not reset the primary evidence accumulator feeding into it. Immediately after (or even during) this resetting, the motor accumulator could resume accumulating, swiftly picking up on new evidence accumulated by the primary accumulator in the interim. In turn, this second wave of motor accumulation can enable the selection of a different response in cases where the new evidence now points towards an alternative choice.

Overall, the LIT has important neurobiological implications as it highlights the need to differentiate between two interrelated but largely separate accumulation processes that are likely to take place in different brain networks. While the former could be independent of sensory and response modality (Heekeren et al., 2006; Philiastides et al., 2011; Filimon et al., 2013; Ploran et al., 2011), the latter would emerge from structures controlling the specific motor effectors involved in implementing the decision (Tosoni et al., 2008; Filimon et al., 2013; Donner et al., 2009), consistent with an embodied cognition model (Cisek & Pastor-Bernier, 2014).

Moreover, we speculate that the all-important interplay between the two processes could be implemented via cortico-basal ganglia circuits controlling the proposed rate of information leakage during motor accumulation. In fact, it is likely that the relevant structures would be comparable to those previously proposed to control the urgency to make a choice by adjusting decision boundaries (Forstmann et al., 2008; Ivanoff et al., 2008; Bogacz et al., 2010; Herz et al., 2017), including work involving oculomotor decisions (Hanks et al., 2014; Jun et al., 2021; Crapse et al., 2018; Stine et al., 2022).

In conclusion, our work provides a novel and biologically plausible alternative to the traditional single evidence accumulation models of choice-RT. Correspondingly, our findings could help revise existing theories of sensorimotor decision making by imposing new mechanistic constraints, while at the same time offering a new benchmark against which the neural systems underlying such decisions can be interrogated.

## Materials and Methods

### Participants

Forty-three right-handed volunteers (18 males) aged between 20 and 48 years (mean = 26 years) participated in the experiment. All participants had normal or corrected-to-normal vision and reported no history of neurological problems. The study was approved by the College of Science and Engineering Ethics Committee at the University of Glasgow (CSE01353), and informed consent was obtained from all participants.

### Stimuli

We used a set of 40 face and 40 car grayscale images (size 500×500 pixels, 8-bits/pixel). Face images were obtained from the Max-Planck Institute of Biological Cybernetics face database Germany (Troje & Bülthoff, 1996). Images of cars were sourced from the web and placed on uniform grey background. We equalized spatial frequency, luminance and contrast across all images. We adjusted the magnitude spectrum of each image to the average magnitude spectrum of all images. The phase spectrum was manipulated to generate noisy images characterized by their percent phase coherence (Dakin et al., 2002). We used two different phase coherence values (25% and 30%) to manipulate the difficulty in the decision. At each difficulty level, we generated multiple frames for each image whereby the spatial distribution of the noise varied across frames, while the overall level of noise remained unchanged. This in turn ensured that when we presented these frames in rapid succession a dynamic stimulus emerged in which relevant parts of the underlying image were revealed sequentially.

### Behavioral Paradigm

Participants performed a visual categorization task by discriminating dynamically updating sequences of either face or car images (Philiastides et al., 2014, 2011). Participants sat at a distance of 75 cm from the presentation screen such that each image sequences was around 6×6° of visual angle. Image sequences were presented in a rapid serial visual presentation fashion at a frame rate of 30 frames per second (i.e., 33.3 ms per frame without gaps) using Presentation (Neurobehavioral Systems, Albany, California). Each trial consisted of a single sequence of noisy images from either a face or car stimulus, at one of the two possible phase coherence levels. We introduced separate Speed and Accuracy instruction blocks (2 blocks per instruction, the order of which was pseudorandomized across participants). To encourage a speed-accuracy tradeoff (SAT), we introduced different response deadlines for the two types of blocks (Speed: 1s, Accuracy: 1.6s).

Participants indicated their choices on a response device using the index finger of their right and left hands. The mapping between face/car choices and left/right index fingers was counterbalanced across participants. Once participants made a response the stimulus was removed from the screen and the trial was terminated. An inter-trial interval then followed and varied randomly in duration in the range 1 – 1.5s. Each block consisted of 160 trials (40 trials per stimulus type and phase coherence level). If participants failed to respond within the allocated time period, a “Too slow!” message appeared on the screen and the trial was marked as a no-choice trial and was excluded from further analysis.

Prior to the main task, we provided training to enable participants to familiarize themselves with the task and allow them time to settle on an appropriate SAT strategy by learning to pace themselves accordingly in each block instruction. Specifically, they trained separately under each of the Speed and Accuracy instruction by completing a total of 80 training trials. During the main experiment, each of the four blocks also contained 10 training trials before the main experimental trials were deployed. Training also ensured that the influence of ongoing learning effects during the main experiments were minimized.

### Prepaid model estimation

To estimate the LIT, a prepaid estimation method for diffusion models of choice RT was used (Mestdagh et al., 2019). First, a prepaid database of time scaled LIT probability distribution functions was created, covering a broad range of model parameters. Because we can adjust the timescale of these distributions at very low computational cost when comparing to data, the parameter degree of freedom responsible for time scaling was not included in the final prepaid grid. The prepaid grid used a uniform distribution for the relative starting point bias 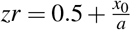 between 0.1 and 0.9 and a uniform distribution for the the inverse leak parameter *λ*^−1^ between 0 and 4. Conditional on each of these 2D grid points, 100 drift speeds *v* wee chosen to span accuracies between 0.001 and 0.999. The boundary separation parameter *a* was determined through time scaling and did not have to be included in the prepaid grid. This was the same grid as used in the original LIT paper. Likewise, a D*M objective function was used as a measure of fit (Verdonck & Tuerlinckx, 2016). A D*M objective function allows the estimation of a choice RT model with an additional, distribution-unspecified non-decision time component in so far that this component is shared across (some) conditions. Using D*M, however, means the time scalar estimator *ŝ* of the original prepaid estimation method (Mestdagh et al., 2019) can no longer be used. Like in the original LIT paper (although not explicitly mentioned in the text), we used a different estimator, which does not depend on the distribution shape of the non-decision component. The appropriate time scalar was estimated by dividing the standard deviation of the observed mean response times *o_c_* of the 4 different stimulus conditions *c*, by its equivalent calculated on the prepaid model distributions *m_c_*:

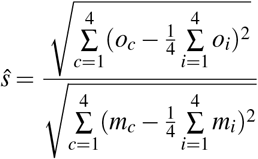

Because the non-decision distribution is the same for all stimulus conditions, its mean cancels out when taking the standard deviation of the response time means of the different stimulus conditions, making this formula independent of that non-decision distribution. Additionally, because D*M is time translation invariant, there is no need to shift the time scaled prepaid distribution before calculating the D*M objective function value in that grid point. The efficacy of this particular prepaid estimation implementation has been extensively demonstrated in the original LIT paper (Verdonck et al., 2021).

### Model selection

#### Approximating the LIT as a reparameterized standard drift diffusion model

In the seminal LIT paper (Verdonck et al., 2021) it was established that any LIT choice RT distribution can be approximated by a standard DDM distribution, transferring the effect of the LIT specific leak parameter *λ* to the rest of the parameters. The impact of the LIT leak on the starting position and non-decision time if analyzed in a standard DDM framework were quantified as follows:

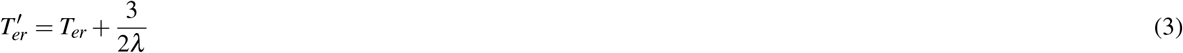

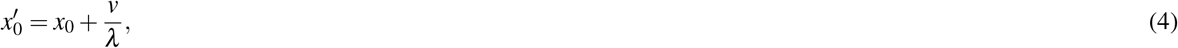

with *x*_0_, *λ* and *T_er_* parameters of the original LIT and 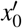 and 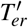 parameters of the resulting DDM.

In the following, we also determine the approximate effect of the LIT leak parameter *λ* on the boundary separation *a′*,which was not yet formally quantified. To do this, we first establish that the deterministic integration of a process of weakly colored noise, as described by Hagan et al. (1989), is equivalent to the LIT as described in Equation 1 applied to a standard DDM model. Hagan et al. describe the following process:

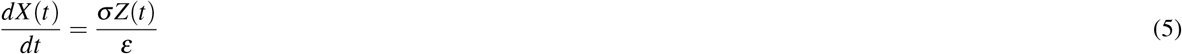

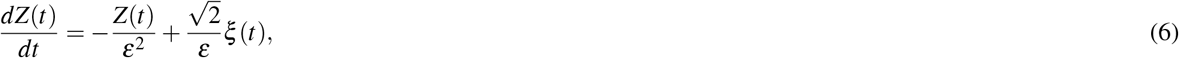

with absorbing boundaries on *X*(*t*) at 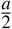 and 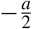. If we take the time derivative of Equation 5, we can substitute 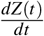 with Equation 6:

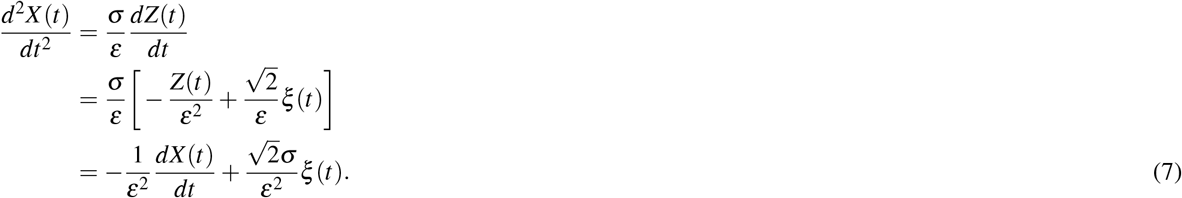

On the other hand, for drift speed *v* = 0, the differential equations for a DDM with a LIT are:

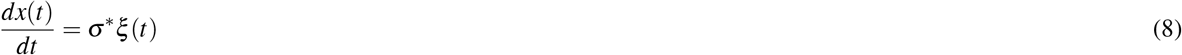

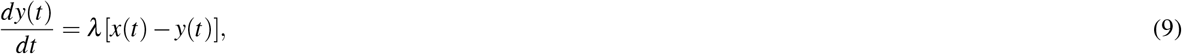

with absorbing boundaries on *y*(*t*) at 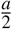 and 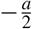. If we take the time derivative of Equation 9, we can substitute 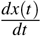 with Equation 8:

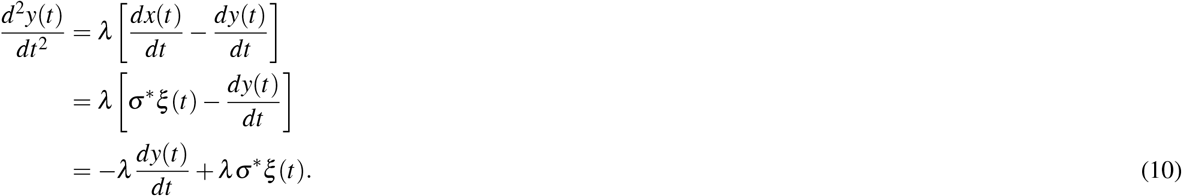

Combining Equation 7 and Equation 10, we can identify the *ε* and *σ* parameters of the problem in Hagan et al. (1989) in terms of DDM-LIT parameters. Comparing the 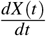 and 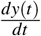 terms, we get:

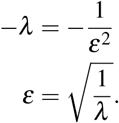

Comparing the *ξ* (*t*) terms, we get

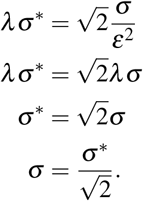

Given the result of Hagan et al. on the increase in boundary separation (Hagan et al., 1989):

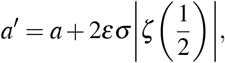

we get

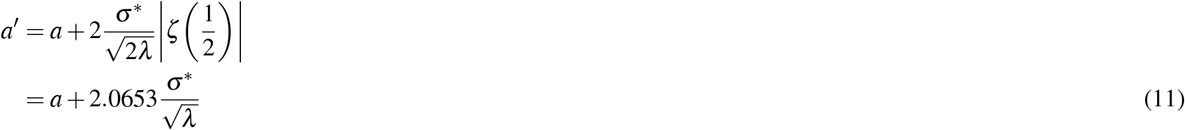

in the LIT context. Drift speed *v* was set to zero in this derivation, but has no impact on it (Hagan et al., 1989). As is standard practice for the DDM, we fix *σ*\* to make the model identifiable. In this paper, we choose *σ*\* = 1.

#### Alternative mechanisms of urgency control

Finally, we can formulate a model that combines three possible mechanisms for controlling urgency: boundary separation *a*, LIT leak *λ* and finally collapse rate *c*:

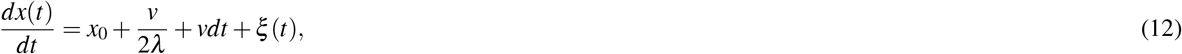

with effective boundary separation 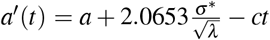 and 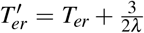. For *c* > 0 This implies a bounded response time distribution, because all responses will occur before time 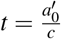, when the collapse is complete.

#### Comparing different mechanisms of urgency control

For each participant separately, we simultaneously fit the choice RT data from the four stimulus conditions and both urgency manipulations. We consider three different models embedded within the framework established in Equation 12. For all models, parameters are assumed to be constant across conditions, except for the drift speed which may change between stimulus conditions (S) and one single additional parameter which may change between urgency conditions (U). The non-decision distribution is also allowed to differ between urgency conditions. As shown in Table 1, the three models under consideration differ only in the parameter that is allowed to change between urgency conditions: The first model assumes the leak parameter (*λ*), the second model assumes the basic boundary separation (*a*) and the third model assumes the collapse rate (*c*) to be responsible for any differences between urgency conditions.

**Table 1.** Constraints of model parameters across conditions, for each of the models under consideration. The constraints are given per model (row): a dot indicates that this parameter is fixed across all conditions, S indicates that the parameter may change between stimulus conditions and U indicates that the parameter may change between urgency conditions.

| model | $T_{er}$ distribution | $a$ | $x_0$ | $v_i$ | $\lambda$ | $c$ |
| --- | --- | --- | --- | --- | --- | --- |
| LIT | U | . | . | S | U | . |
| fixed boundary | U | U | . | S | . | . |
| collapsing bounds | U | . | . | S | . | U |

The probability density functions are calculated using the grid-based fast-dm approach (Voss & Voss, 2008), but recoded for GPU for increased performance. Parameters are estimated using a D*M objective function (Verdonck & Tuerlinckx, 2016). The objective function is minimized using the differential evolution approach (Storn & Price, 1997). As all models under consideration have the same number of parameters, we can straight up compare fits for model selection purposes.

### Parameters used for prediction simulations

All simulations are done using a normal Euler-Maruyama algorithm with a dt accuracy of 1 ms. The simulations needed to create Figure 3 use the parameters given in Table 2 for each condition. Additionally, all conditions use a starting position *x*_0_ = 0. For each condition, 100000 trials were run. Finally, before calculating the correlation-RT curves, an amount of normally distributed noise (standard deviation 0.015) was superimposed on the mean signals to simulate measurement noise and residues from other processes.

**Table 2.**
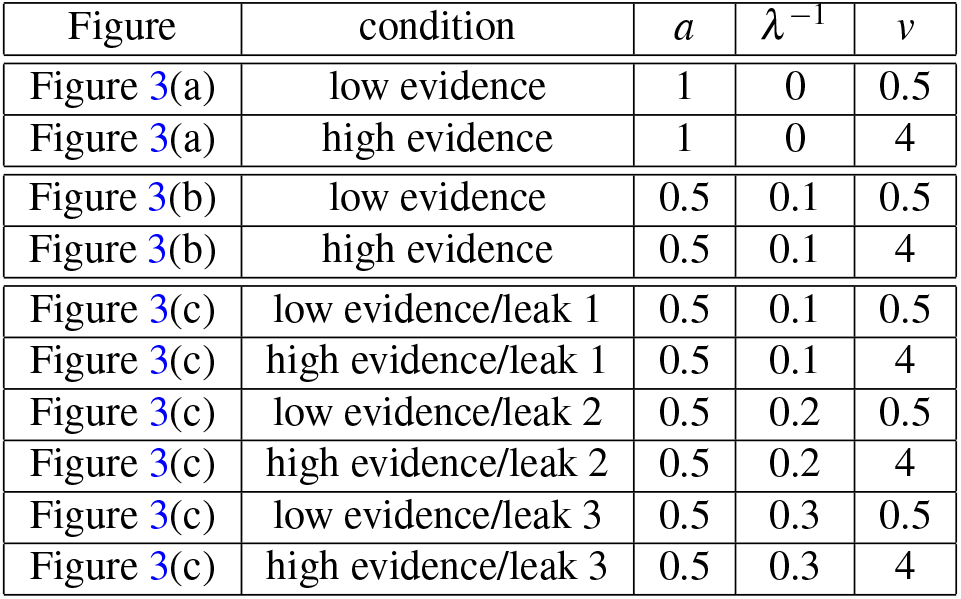
Prediction simulation parameters.

### EEG data acquisition

We collected EEG data inside an electrostatically shielded booth using a 64-channel EEG amplifier system (BrainAmps MR-Plus, Brain Products, Germany) and recorded using Brain Vision Recorder (BVR; Version 1.10, Brain Products, Germany) with a 1kHz sampling rate and an analogue bandpass filter of 0.016–250 Hz. The EEG cap consisted of 64 Ag/AgCl passive electrodes (Brain Products, Germany) positioned according to the international 10–20 system of electrode positioning, with chin ground and left mastoid reference electrode. All input impedances were kept below 20 kΩ. Experimental event codes and behavioral responses were also synchronized with the EEG data and collected using the BVR software.

### EEG pre-processing

We pre-processed the EEG signals offline using MATLAB (MathWorks, Natick, Massachusetts). Specifically, we applied a bandpass (4th order Butterworth) filter between 0.5 and 40 Hz. The filter was applied non-causally to avoid distortions caused by phase delays (using MATLAB ‘filtfilt’). We then removed eye-blink artifacts using a principal component analysis (PCA) approach. Specifically, prior to the main experiment, we asked participants to complete an eye movement calibration task during which they were instructed to blink repeatedly upon the appearance of a fixation cross in the centre of the screen, while we collected EEG data. This enabled us to determine linear EEG sensor weightings corresponding to eye blinks using PCA such that these components were projected onto the broadband data from the main task and subtracted out (Philiastides et al., 2006; Gherman & Philiastides, 2018). Finally, we re-referenced the EEG data to the average of all channels.

### EEG analysis

#### Estimating the evidence accumulation signal

Starting from the assumption that any meaningful evidence accumulation signal should be able to discriminate between stimulus conditions of high versus low sensory evidence, we estimate it as the linear combination of signal components, *y*(*t*), that is best able to discriminate between these conditions:

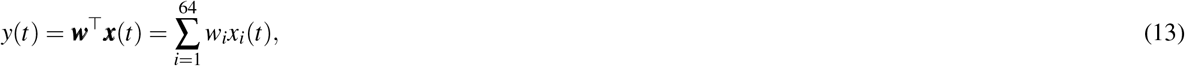

with ***w*** the sensor weights (spatial filter) resulting in a maximally discriminating *y*(*t*).

We perform this linear discriminant analysis across response-locked time points, by using a sliding window approach. Specifically, we define time windows of 50 ms duration and shift the window centre with 25 ms increments in a time interval ranging from 575 ms before to 175 ms after the response. For each sensor, we then time-average the samples within each 50 ms window. Using these time-averaged channel values, the sensor weights that maximally discriminate between trials with different levels of sensory evidence, can be found using the Fisher discriminant (Stork et al., 2001):

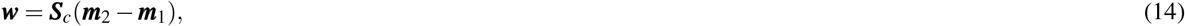

with

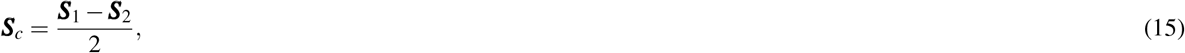

where ***m***_i_ is the estimated mean and ***S***_i_ is the estimated covariance matrix of the time-averaged channel values of the trials of condition *i =* 1,2 (i.e., low and high sensory evidence trials). We quantify the performance of this classifier for each time window by using the area under a receiver operator characteristic (ROC) curve (Green et al., 1966), referred to as *Az*, with a leave-one-out cross-validation approach (Stork et al., 2001). Scalp topographies are calculated by projecting the raw signal of each sensor on the already established evidence accumulation signal vector ***y*** (50×1), evaluated in the 50ms window centered around the response:

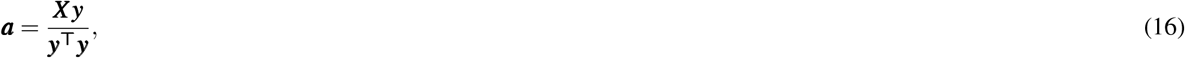

with ***X*** (64×50) a matrix composed of 64 channel rows covering 50ms of signal. Vector ***a*** describes the electrical coupling between the individual sensors and the evidence accumulation signal.

#### Estimating the motor accumulation signal

The fundamental dynamical assumption of the LIT is comprised in differential Equation 1, where it is stated that small changes in the motor accumulation signal are proportionate to the momentary value of the evidence accumulation signal. Based on this we would expect, for each time *t* of each separate trial, the momentary slope of the motor accumulation signal to be linearly proportional to the momentary value of the evidence accumulation signal. This proportionality should result in a correlation between all trial-wise momentary slopes from the motor accumulation signals and their corresponding momentary evidence accumulation signal values. In what follows we try to find those sensor weights and resulting aggregate motor accumulation signal, that maximizes this correlation in the data.

For each participant separately, we calculate a number of slope snapshots. Each slope snapshot is calculated based on the raw signals of a specific trial and pertains to individual 50 ms time windows within that trial. It is a vector containing the slopes (linear regression coefficient) of the signal at every sensor (here the number of sensors is 64). For each trial, we calculate a 50 ms window slope snapshot every 25 ms with window centres ranging from 150 ms to 50 ms before the response. The total number of slope snapshots, which is the product of the number of trials under consideration and the number of time windows for each trial, is defined as *K*. From the slope snapshots *S_sk_* (64 x *K*), we estimate the sensor weights *w_s_* that produce a spatially aggregated slope that is maximally correlated with previously calculated evidence accumulation signals *e_k_* (1 x *K*), using the following formula:

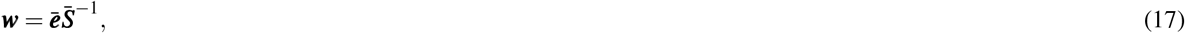

with ***ē*** defined by

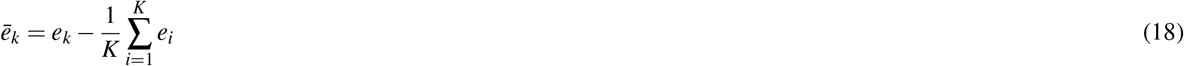

and 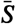 defined by

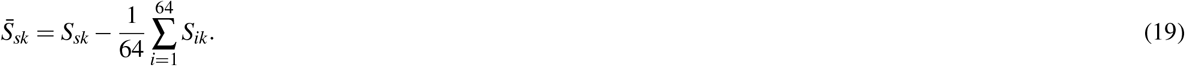

In this analysis, we only use the high sensory evidence trials, to enhance potential left-right hemisphere differences.

## Data and code availability

All data (behavioral and EEG) will be made freely available prior to the publication of the manuscript. Code can be obtained from the authors upon request.

## Acknowledgements

This work was supported by the European Research Council (865003; MGP) and the Economic and Social Research Council (ES/L012995/1; MGP), the Research Fund of KU Leuven (C14/19/054; SV) and FWO (G074219N; SV). We thank Bharti Gupta for assistance with data collection.

## Author contributions

M.P. and S.V. conceived the experiment, M.P. conducted the experiment, S.V. developed the new modeling and validation approaches, T.L. developed the theoretical work for the extension of the LIT approximation, S.V. analysed the data, M.P. and S.V. interpreted the results. All authors wrote the manuscript.

